# Single-cell chromatin interactions reveal regulatory hubs in dynamic compartmentalized domains

**DOI:** 10.1101/307405

**Authors:** A. Marieke Oudelaar, James O.J. Davies, Lars L.P. Hanssen, Jelena M. Telenius, Ron Schwessinger, Yu Liu, Jill M. Brown, Damien J. Downes, Andrea M. Chiariello, Simona Bianco, Mario Nicodemi, Veronica J. Buckle, Job Dekker, Douglas R. Higgs, Jim R. Hughes

## Abstract

The promoters of mammalian genes are commonly regulated by multiple distal enhancers, which physically interact within discrete chromatin domains. How such domains form and how the regulatory elements within them interact within single cells is not understood. To address this we developed Tri-C, a new Chromosome Conformation Capture (3C) approach to identify concurrent chromatin interactions at individual alleles within single cells. The heterogeneity of interactions observed between such cells shows that CTCF-mediated formation of chromatin domains and interactions within them are dynamic processes. Importantly, our analyses reveal higher-order structures involving simultaneous interactions between multiple enhancers and promoters within individual cells. This provides a structural basis for understanding how multiple *cis*-elements act together to establish robust regulation of gene expression.

## Introduction

Precise spatial and temporal patterns of gene expression during development and differentiation are controlled by *cis*-regulatory elements including promoters, enhancers and boundary elements. The interaction and activity of these elements depend on their structural organization within the nucleus (1). To date the relationship between structure and function has mainly been analyzed in populations of cells. The globin loci, which provide ideal models to elucidate the general principles of mammalian gene regulation, have been extensively studied in this way. For example, we have previously shown that the active mouse α-globin cluster and its regulatory elements are located in a decompacted ~70 kb chromatin domain that forms early in erythroid differentiation and is flanked by CTCF-binding sites (2–4). Within this domain, the α-globin genes interact with five enhancer elements, which cooperate in an additive manner to upregulate gene expression (5). Such activity of multiple enhancer elements has been frequently observed in a wide variety of mammalian gene loci and is thought to impart robustness to patterns of gene expression underlying cell fate decisions (6).

It is now known that analyzing populations of cells may not reveal the true mechanisms underlying biological processes that are revealed by single cell analysis. At present, we do not know how promoters, enhancers and boundary elements physically interact as genes are switched on in single cells. To address this important question, we developed Tri-C, a new Chromosome Conformation Capture (3C) technique, which can identify concurrent chromosomal interactions at individual alleles, and consequently provides information derived from single cells. We combined conventional 3C and Tri-C experiments to perform in-depth characterization of the structural architecture of the murine globin loci in single cells.

Our analyses of the single-cell conformations of the domains containing the globin loci show that interactions between boundary elements are heterogeneous, implying that they are dynamic rather than fixed, thus providing support for a loop extrusion mechanism contributing to the formation of chromatin domains. Furthermore, we observe higher-order complexes within these domains in which multiple enhancers and promoters interact within individual nuclei. This shows that more than one enhancer frequently interacts simultaneously with individual promoters within a single cell. This is consistent with previous observations showing that fully regulated expression of the murine globin genes (5, 7) and many other genes (6) depends on the presence of more than one enhancer element. The formation of such enhancer-promoter complexes also explains the function of apparent redundant enhancer elements (8, 9), which could play a role in the formation of robust structures required for assembly of the transcriptional machinery at gene promoters.

## Results

### Activation of the murine globin loci is associated with the formation of strongly compartmentalized domains in which enhancers and promoters interact

To characterize the large-scale conformations of the murine globin loci, we performed Hi-C in primary mouse erythroid cells, in which the globin genes are highly expressed, and compared this to an equivalent Hi-C dataset from mouse embryonic stem (ES) cells (10), in which the globin loci are silent (Supplementary Figure 1). This comparison shows that the chromatin regions containing the globin clusters form strong, tissue-specific, self-interacting domains in erythroid cells, but not in ES cells. To characterize the interactions within these domains at higher resolution, we performed multiplexed, high-resolution Capture-C experiments in erythroid and ES cells from ∼45 viewpoints, which were closely spaced across the loci and included all known regulatory elements. Figure 1 shows interaction profiles in the α-globin domain from the viewpoint of the promoters, the strongest enhancer element (R2), and the CTCF boundary elements on either side of the domain. These profiles show strong reciprocal interactions between the α-globin promoters and all its enhancer elements in erythroid cells. The flanking CTCF-binding elements do not participate in these enhancer-promoter interactions, but form diffuse interactions with the chromatin bound by CTCF on either side of the domain, spanning contiguous regions of ∼50 kb. The nature of these interaction patterns is highlighted in the contact matrices derived from the viewpoints tiled across a 300 kb region containing the locus (Figure 2).

**Figure 1:**
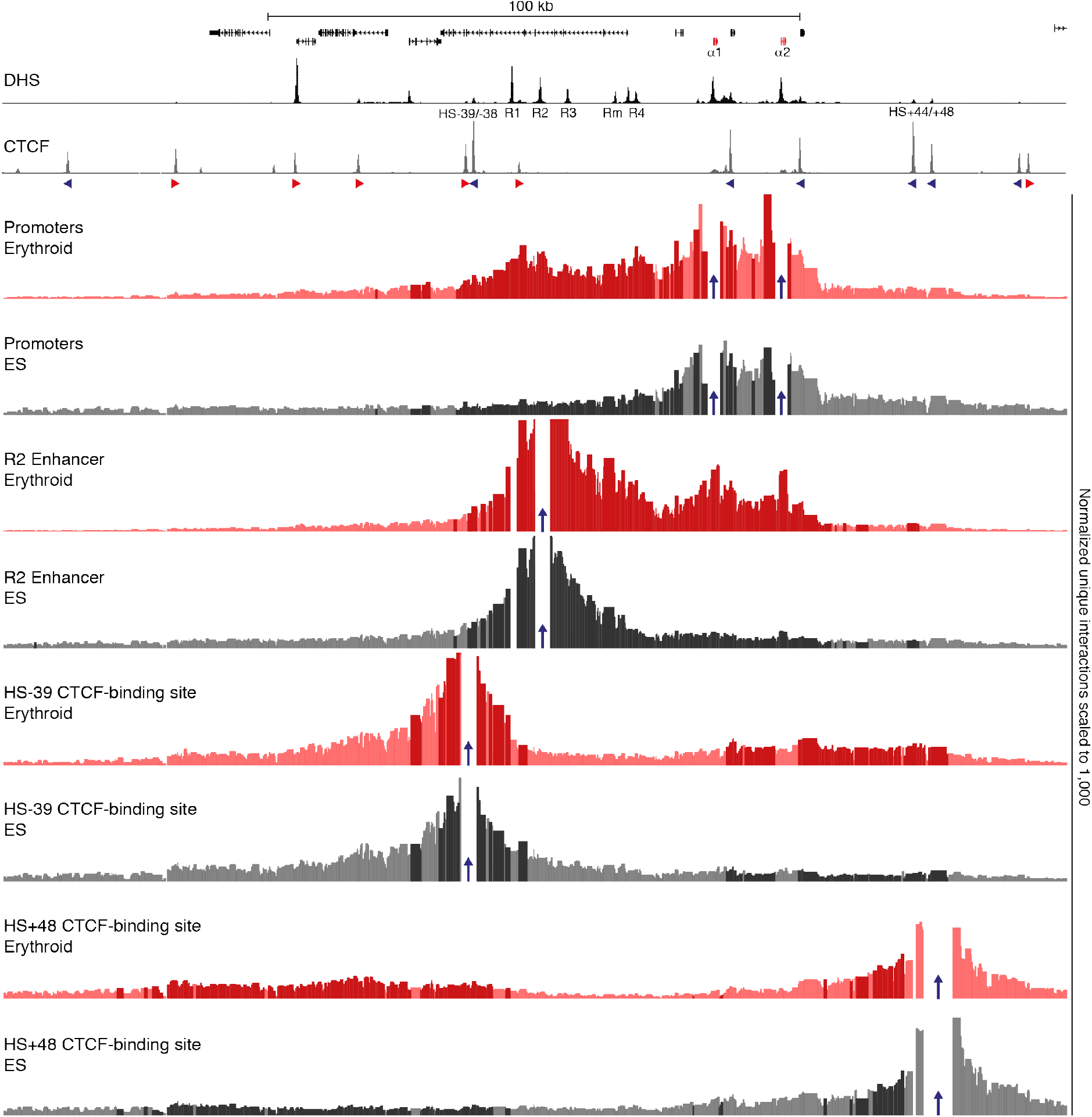
Characterization of the interaction landscape of the regulatory elements of the α-globin locus. High-resolution Capture-C interaction profiles of the α-globin locus from the viewpoints (indicated by blue arrows) of the α-globin promoters, the R2 enhancer, and CTCF-binding sites HS-39 and HS+48 in erythroid (red) and ES (grey) cells. Profiles represent the mean number of normalized unique interactions per restriction fragment from three replicates. Significantly different interactions between erythroid and ES cells are highlighted in bold colors. Gene annotation (α-globin genes highlighted in red), erythroid DNaseI Hypersensitive Sites (DHS) and CTCF occupancy are shown at the top, with arrows indicating the orientation of the CTCF-binding motifs. Coordinates (mm9): chr11:32,050,000-32,250,000.

**Figure 2:**
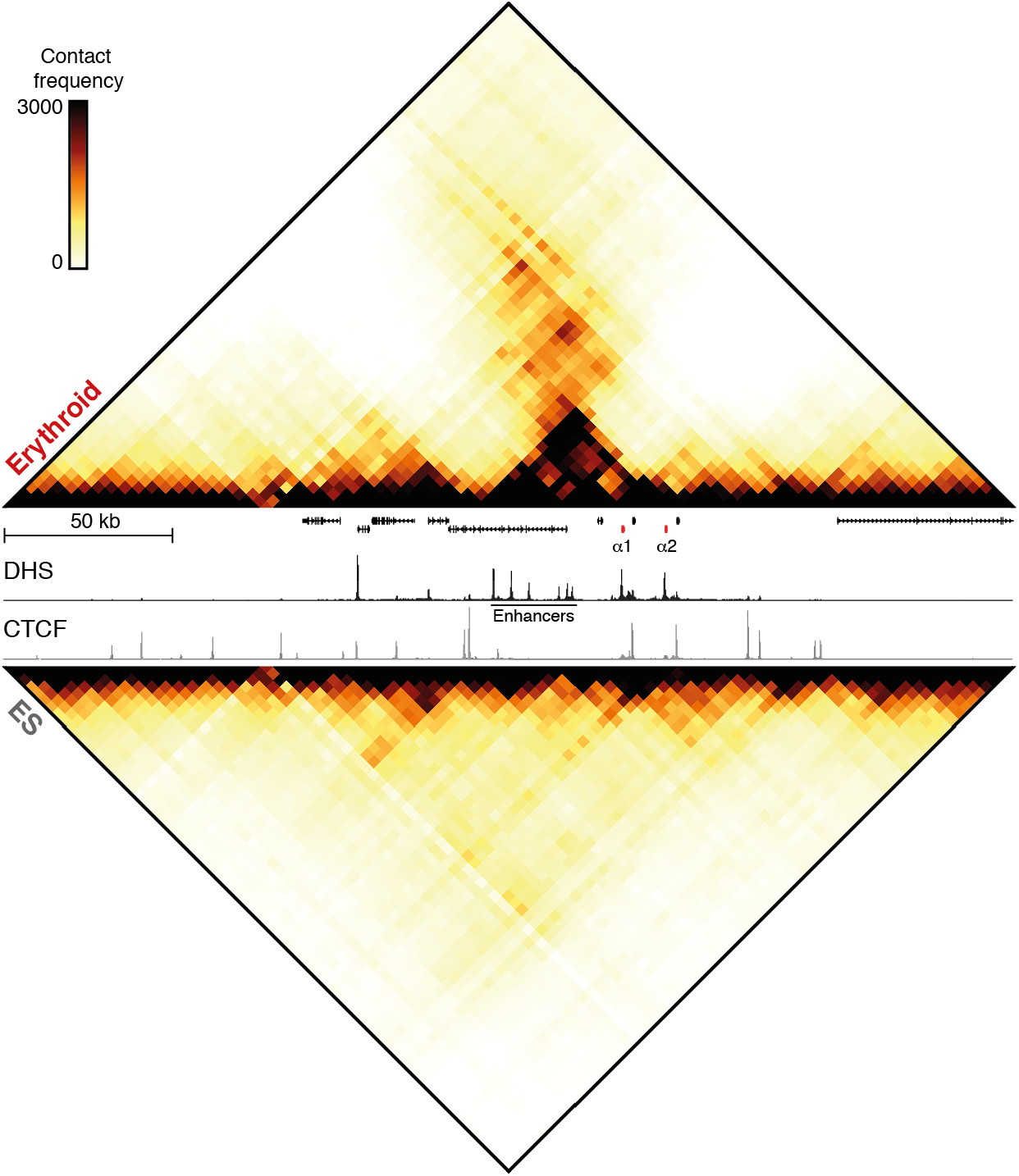
Structural conformation of the active and inactive α-globin locus. Contact matrices (4 kb resolution) of the α-globin locus derived from Capture-C experiments with viewpoints closely spaced across the domain in erythroid (top) and ES (bottom) cells. Contact frequencies represent the mean number of normalized unique interactions from three replicates. Gene annotation (α-globin genes highlighted in red), erythroid DNaseI Hypersensitive Sites (DHS) and CTCF occupancy are shown in the middle. Coordinates (mm9): chr11:32,000,000-32,300,000.

Although the contacts between the enhancers and promoters are more discrete compared to those between the CTCF-binding sites, interaction frequencies are significantly enriched throughout the domain in erythroid cells. Importantly, this is not consistent with a model in which enhancers and promoters simply form stable, distinct loops (1, 11). Rather, these interactions are more readily interpreted as a compartmentalized domain, in which, at some point, every region of chromatin interacts with every other, and preferred, transiently stabilized structures are formed between regulatory elements. Analysis of the β-globin locus reveals a similar interaction landscape (Supplementary Figures 2, 3).

Our analysis by Capture-C suggests that all enhancers and promoters in the globin loci can form interactions. However, as these data are derived from pair-wise contacts in populations of cells, they reflect multiple dynamic conformations that obscure specific higher-order structures associated with productive interactions between enhancers and promoters in individual cells. In particular, it is not clear whether *cis*-interactions between multiple enhancers and promoters occur simultaneously in a single cell nucleus, or if these elements compete for the formation of exclusive interactions. Therefore, it remains unclear how these regulatory elements interact to ensure robust regulation of gene expression.

### Tri-C detects multi-way interactions with unprecedented sensitivity

Questions regarding the structural interaction between regulatory elements can be addressed by analyzing interactions between these elements in individual cells. Though single-cell Hi-C (12–14) and Genome Architecture Mapping (15) have provided insights into chromosomal structures in single cells at large scales, the resolution of these techniques (100 kb and 30 kb, respectively) is insufficient to be informative at the level of individual regulatory elements. To overcome these limitations, we explored a different strategy to analyze chromosomal structures in single cells. Because *in situ* 3C libraries contain long DNA concatemers in which neighboring fragments represent chromatin regions that were in close proximity in individual nuclei, single-cell chromatin conformations can be derived from population-based assays in which several neighboring fragments in the 3C concatemer are identified simultaneously. However, the proportion of reads that contain multiple interacting fragments in conventional 3C techniques is very low (<2%) (16–19) and despite advances in efficiency in a recent 3way-4C approach (20), resolution and sensitivity of current techniques are insufficient for robust detection of higher-order structures formed by individual regulatory elements in single cells.

To analyze such structures at high resolution, we developed Tri-C, a new 3C method that can identify multi-way interactions with viewpoints of interest with unprecedented sensitivity (Figure 3a). For efficient detection of multiple ligation junctions between interacting fragments, Tri-C libraries are generated using an enzyme selected to create relatively small DNA fragments (∼200 bp) for the viewpoints of interest. We chose the restriction enzyme NlaIII, which has a recognition sequence of four bp and therefore cuts on average every 256 bp. Sonication of these libraries to ∼450 bp generates fragments of which about ∼50% contain multiple ligation junctions. Using highly optimized oligonucleotide capture-mediated enrichment for viewpoints of interest, millions of multi-way contacts can be identified with Illumina sequencing, generating 3C profiles at unprecedented depth (Figure 3b). Importantly, Tri-C allows for multiplexing both viewpoints and samples, enabling analyses of multiple genomic regions and cell types of interest in a single experiment. Because Illumina sequencing provides accurate identification of the random sonication ends of the reads, these can be used as unique molecular identifiers to filter out PCR duplicates, allowing for quantitative analysis of the detected multi-way interactions.

**Figure 3:**
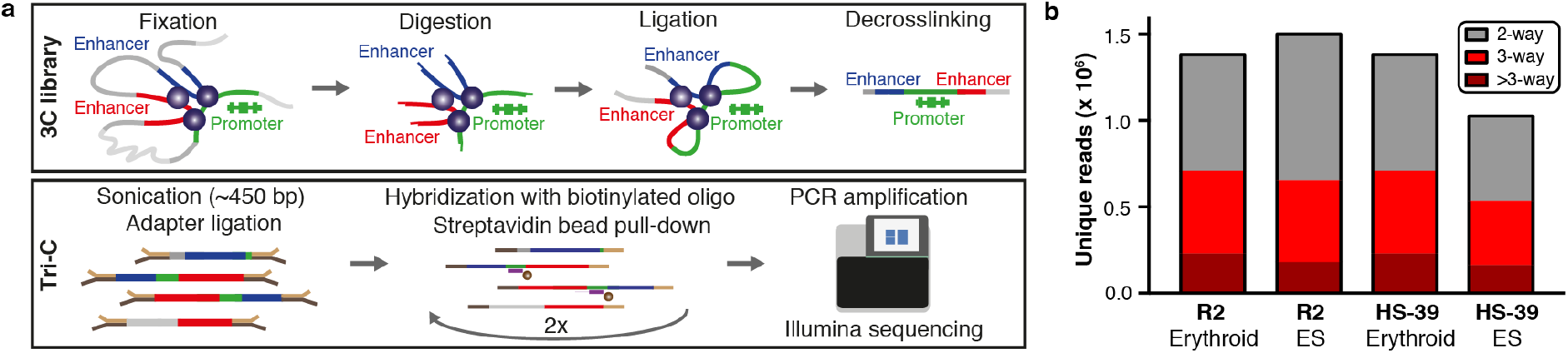
Overview of the experimental procedure and data output of Tri-C. **(a)** Overview of Tri-C. **(b)** Number of unique reads containing pair-wise and multi-way interactions generated by Tri-C for viewpoints in the α-globin locus.

To validate that Tri-C detects reliable multi-way interactions in individual cells, we performed several additional experiments and analyses. First, to confirm that capturing multiple ligation junctions simultaneously did not introduce a bias in the detected interactions, we compared the pair-wise interactions derived from Tri-C to Capture-C interaction profiles from the same viewpoint (Supplementary Figure 4). Next, we validated the detected multi-way interactions by long-read Nanopore sequencing of PCR-enriched ligation events (Supplementary Figure 5). Finally, to confirm that Tri-C interactions represent single-cell chromosome conformations, we show that the detected multi-way interactions are allele-specific (Supplementary Figure 6).

**Figure 4:**
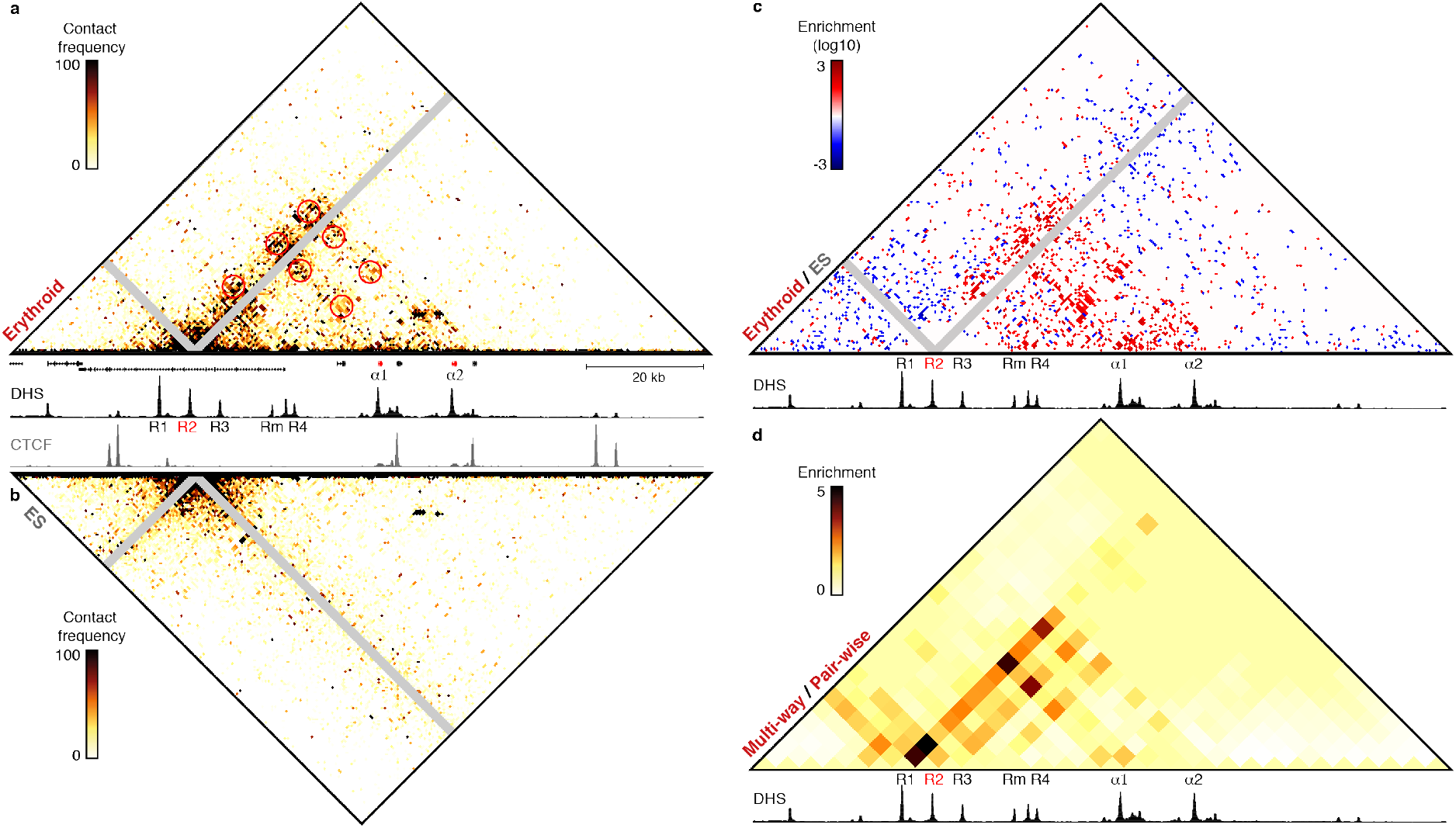
Analysis of multi-way interactions between enhancers and promoters in the α-globin locus. Tri-C data (500 bp resolution) showing multi-way chromatin interactions with the R2 enhancer in the α-globin locus. Gene annotation (α-globin genes highlighted in red), erythroid DNasel Hypersensitive Sites (DHS) and/or CTCF-binding sites are shown at the bottom of the matrices, **(a)** Contact matrix showing normalized multi-way interactions in three replicates of erythroid cells. Proximity contacts around R2 are excluded (grey diagonal) and specific enrichments are highlighted (red circles). **(b)** Contact matrix showing normalized multi-way interactions in three replicates of ES cells. Proximity contacts around R2 are excluded (grey diagonal). **(c)** Contact matrix highlighting interactions that are >20-fold enriched in erythroid (red) or ES (blue) cells. Proximity contacts around R2 are excluded (grey diagonal). **(d)** Contact matrix (4 kb resolution) showing multi-way interactions after correcting for the pair-wise contact frequencies derived from the multiplexed Capture-C data. Coordinates (mm9): chr11:32,120,000-32,240,000.

**Figure 5:**
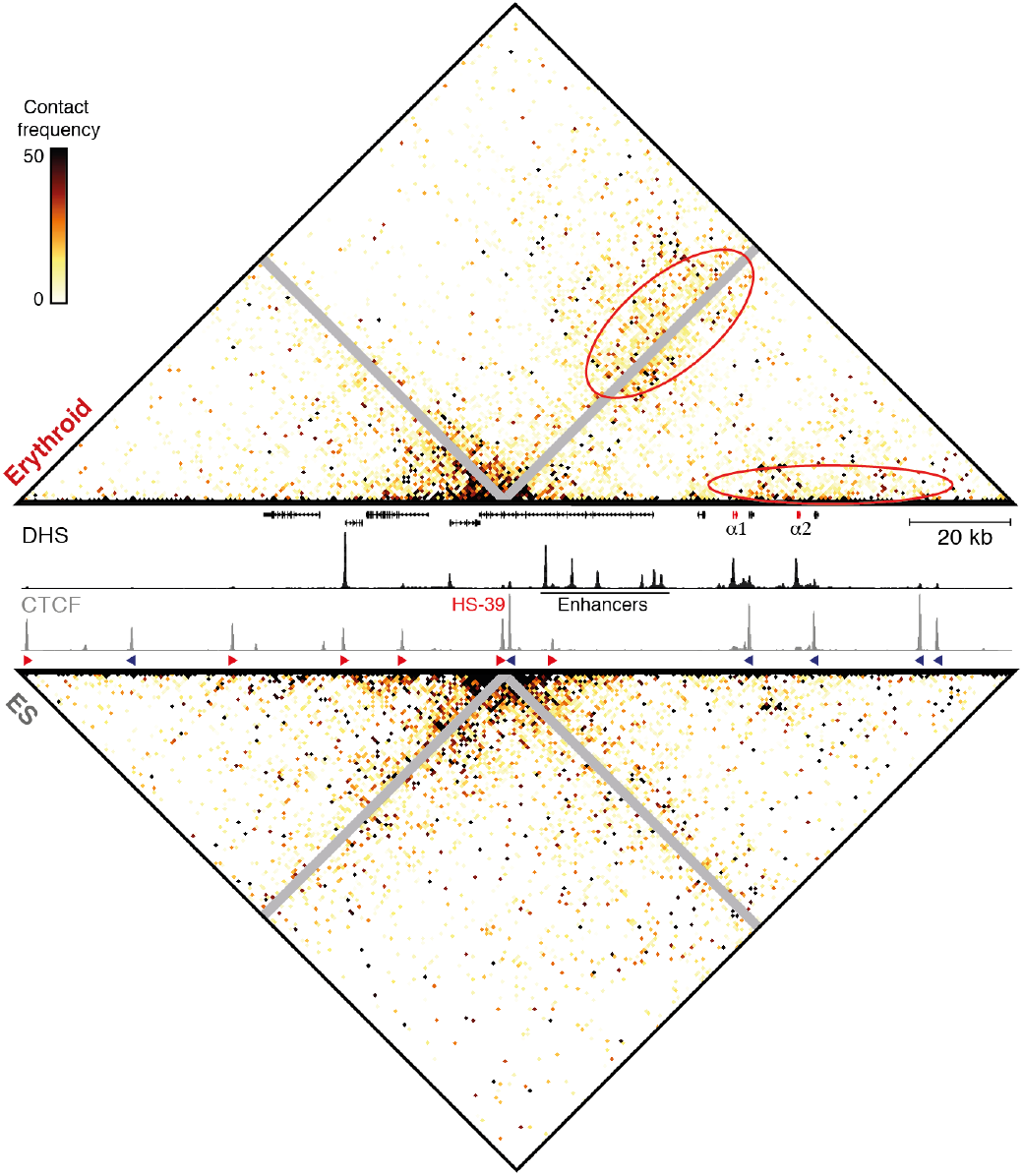
Analysis of multi-way interactions between CTCF-binding sites in the α-globin locus. Tri-C contact matrices (500 bp resolution) showing normalized multi-way chromatin interactions with CTCF-binding site HS-39 in the α-globin locus in three replicates of erythroid (top) and ES (bottom) cells. Gene annotation (α-globin genes highlighted in red), erythroid DNaseI Hypersensitive Sites (DHS) and CTCF-binding sites are shown in the middle, with arrows indicating the orientation of the CTCF-binding motifs. Proximity contacts around HS-39 are excluded (grey diagonal) and specific enrichments in erythroid cells are highlighted (red ovals). Coordinates (mm9): chr11:32,040,000-32,240,000.

### Multiple enhancers and promoters form higher-order chromatin structures in which they interact simultaneously

We used Tri-C to analyze the higher-order structures around regulatory elements in the globin loci in erythroid and ES cells. To visualize these 3D structures, we represent the multiway interactions in contact matrices in which we excluded the viewpoint of interest and plotted the frequencies with which two elements were captured simultaneously with this viewpoint. Mutually exclusive contacts between elements appear as depletions at the intersections between such elements in the contact matrix, whereas preferential simultaneous interactions in higher-order structures are visible as enrichments at these foci.

Analysis from the R2 viewpoint, the strongest α-globin enhancer (5), shows clear enrichments at the intersections with the other enhancers (R1, R3, Rm and R4) and promoters in erythroid cells (Figure 4a). These interactions are not observed in ES cells (Figure 4b) and direct comparisons show over 100-fold enrichment for the multiple enhancer-promoter interactions in erythroid cells (Figure 4c), demonstrating that these do not simply reflect genomic proximity. These patterns and enrichments are highly reproducible between biological replicates (Supplementary Figure 7). Moreover, the interactions are enriched compared to the contact distribution detected by the multiplexed Capture-C experiments (Figure 4d), highlighting that these higher-order structures cannot be predicted by pair-wise 3C data. Tri-C analysis of HS2, one of the strongest β-globin enhancers (7), shows very similar structures (Supplementary Figure 8).

These analyses demonstrate that multiple enhancers and promoters form higher-order structures in which they simultaneously interact together to switch genes on. Of interest, the promoter of the housekeeping gene *Nprl3*, which is six-fold upregulated in erythroid cells (2), is included in the complexes formed with the α-globin enhancers and promoters, highlighting that there is no competition between genes for mutually exclusive interactions with enhancers.

### CTCF-binding sites form dynamic interactions supportive of a loop extrusion mechanism underlying boundary formation

We have previously shown that CTCF-binding sites flanking the α-globin locus contribute to the formation of a domain that delimits the region of chromatin within which the observed complexes between enhancers and promoters can be formed (2). However, the processes underlying the formation of such chromatin domains and their contribution to enhancer-promoter specificity remain unclear. Based on Hi-C data, it has been suggested that CTCF-binding sites located at domain boundaries form specific loops and that multiple CTCF-binding sites might form multi-anchored and/or nested structures (21). Our Capture-C data show that the functionally validated CTCF-binding site HS-39 (2) located upstream of the a-globin cluster predominantly interacts with a region downstream of the domain, which also contains many CTCF-binding sites (Figures 1 and 2). However, the pattern of interactions is very broad compared to the interactions between enhancers and promoters and it is unclear whether these interactions represent stable loops between multiple CTCF-binding sites or represent a more dynamic mechanism by which domain boundaries are formed.

To resolve these structures within individual nuclei, we performed Tri-C analysis from the viewpoint of the HS-39 CTCF-binding site (Figure 5). Consistent with the pair-wise interaction data, the multi-way interactions are preferentially formed with the region located downstream of the α-globin domain containing many CTCF-binding sites. A model in which CTCF boundaries are formed by stable, multi-anchored loops would result in specific enrichments of multi-way contacts between these CTCF-binding sites. However, the Tri-C contact matrix in erythroid cells shows a diffuse enrichment with the entire region on the other side of the α-globin domain. This diffuse enrichment is confined to the viewpoint diagonal and the base of the matrix, which indicates that the only simultaneous interactions we observe with the HS-39 CTCF-binding site are proximal to HS-39 itself or to its interacting partner. This shows that CTCF-binding sites do not form specific higher-order structures. Rather, the diffuse pattern indicates that HS-39 forms dynamic interactions with the entire region flanking the opposite side of the α-globin domain. The spread of interactions suggests that conformations seen in single cells represent snapshots of a dynamic scanning process throughout this chromatin region.

Tri-C analysis from the viewpoint of the 3’HS1 CTCF-binding site in the β-globin locus shows a similar pattern (Supplementary Figure 9). Our data are therefore not consistent with the formation of stable loops between CTCF-binding sites at domain boundaries. Rather, the Tri-C interaction patterns provide support for a loop extrusion mechanism, in which the formation of chromatin domains is mediated by protein complexes, likely involving Cohesin, that translocate along chromosomes, bringing every region in contact with each other (22–24). Continuous scanning across chromatin regions and transient stalling of these protein complexes at CTCF boundary elements explains the enrichment over a broad region of chromatin containing many CTCF-binding sites. In contrast to these preferential interactions with the opposite boundary region in erythroid cells, the multi-way interactions of CTCF-binding sites around the globin clusters in ES cells are more symmetrical along the genome. However, the patterns of interactions, with broad enrichments along the diagonal and the base of the matrix, are similar, indicating that loop extrusion is a general mechanism contributing to chromosome organization in all cell types.

## Discussion

Formation of chromatin domains by the proposed loop extrusion mechanism could explain many features of chromosome organization (22–24). However, the process of loop extrusion and the resulting dynamic chromatin structures have not been observed directly, and current evidence is derived from polymer model predictions (23, 24) and perturbations of specific components of the loop extrusion machinery (25–27). Here, we show for the first time that high-resolution chromatin structures in single cells do not support a model of stable loops between CTCF-binding sites, but indicate a dynamic mechanism such as loop extrusion underlying the formation of chromatin domains. Importantly, the diffuse enrichment patterns we observe are indicative of transient CTCF-binding site interactions, which is in agreement with the kinetics of CTCF binding to the genome (28).

The preferential interactions between CTCF-binding sites at opposite ends of the globin domains in erythroid cells and the more symmetrical patterns in ES cells explain the formation of tissue-specific domains in erythroid cells (Figure 5 and Supplementary Figure 9). We have shown before that CTCF occupancy is similar in both cell types (2). These different structures therefore likely reflect differences in the processivity of loop extruding factors such as Cohesin, which could result from tissue-specific recruitment locations and/or extrusion rates. It has been shown that Mediator, a transcriptional coactivator bound at active enhancers and promoters, forms complexes with Cohesin and the Cohesin-loading factor Nipbl (29), and we have previously observed binding of Cohesin and Mediator at the α-globin enhancers in erythroid cells (2, 5). The erythroid-specific formation of the globin domains could therefore be explained by a loop extrusion model in which Cohesin is recruited to the genome at active enhancers and/or promoters. Recent findings have suggested that Cohesin translocation is stimulated by Nipbl, suggesting that Nipbl abundance determines extrusion rates (30). Differences in Nipbl distribution could therefore also contribute to tissue-specific interaction patterns, though this remains to be further explored. Within the dynamic chromatin domains containing the globin gene clusters, we identify higher-order structures in which multiple enhancers and promoters interact simultaneously in individual cells. These complex, tissue-specific structures, cannot be explained by CTCF/Cohesin-mediated loop extrusion alone and indicate other, independent mechanisms contributing to chromosome architecture. This is consistent with polymer models (31) and recent studies in which Cohesin binding was perturbed, but interactions between enhancers and promoters still occurred, though more promiscuous (25, 26). Importantly, contacts between these elements do not represent stable, exclusive enhancer-promoter loops (11), but preferred interactions within dynamic compartmentalized domains, in agreement with polymer model predictions (32). Our data could therefore be explained by a loop extrusion mechanism that brings regulatory elements into close proximity and enables subsequent formation of more complex, stabilized structures. This is likely mediated by multi-protein complexes and could contribute to or result from the formation of phase-separated assemblies of components of the transcriptional machinery (33) (Figure 6).

**Figure 6:**
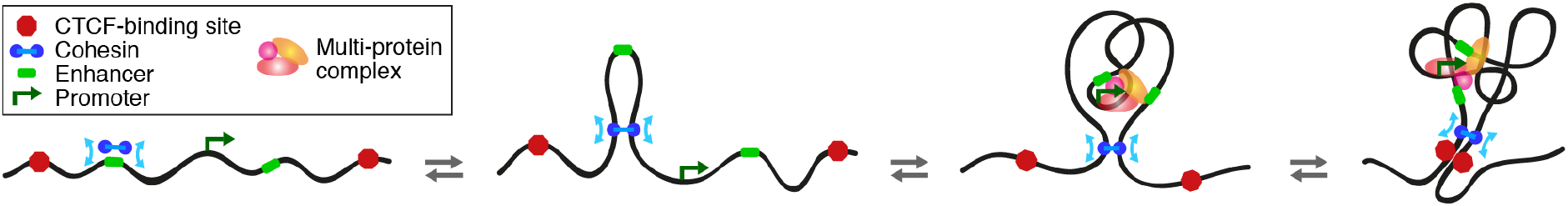
Graphical summary. Our data are supportive of a loop extrusion mechanism contributing to the formation of compartmentalized chromatin domains. Complex higher-order structures, in which multiple enhancers and promoters interact, are formed within these domains by distinct tissue-specific mechanisms, likely involving the formation of phase-separated protein assemblies of the transcriptional machinery.

The formation of such complexes in which multiple enhancers simultaneously interact with the genes they regulate indicates cooperation rather than competition between enhancer elements. This is consistent with the observed additive effects of individual enhancers at the globin (5, 7) and many other genes (6). Importantly, we have previously shown that no single enhancer element in the α-globin locus is critical for the formation of the chromatin structures associated with active α-globin transcription (3, 5). The formation of complexes in which multiple enhancers and promoters interact simultaneously therefore provides a structural basis for the observed functional cooperativity and also suggests a role for apparent redundant enhancer elements (8, 9). Such ‘shadow’ enhancers could have a structural function in forming and maintaining effective platforms for assembly of the transcriptional machinery and ensure the formation of robust complexes, even in the context of mutations or deletions in other enhancer elements.

This highlights that Tri-C analyses not only contribute to our fundamental understanding of the relationship between genome structure and function, but are also a valuable tool to interpret how genetic variations can disrupt complex chromatin structures and cause misregulation of gene expression and disease.

## Methods

### Cells

Primary murine ter119+ erythroid cells were obtained from spleens of female C57BL/6 mice treated with phenylhydrazine as previously described (34). Mouse embryonic stem (ES) cells (129/Ola) were derived from mice at embryonic day 14 and cultured and harvested as previously described (34).

### Hi-C – Experimental procedure

Hi-C in primary murine ter119+ erythroid cells was performed as previously described (35). An equivalent dataset in mouse ES cells (Cast/129) was used for comparative analyses (10).

### Hi-C – Data analysis

Hi-C data were analyzed using the HiC-Pro pipeline (36). Reads were aligned to the mm9 reference genome using Bowtie2, with minor modifications to the recommended options (erythroid: -k 3 –score-min L,-0.6,-0.2; ES: -k 3 –score-min L,-0.6,-0.6) to allow for multimapping in the duplicated regions in the globin loci and better alignment of the ES data in the β-globin region, which contains many SNPs in the Cast/129 strain compared to the mm9 reference.

TADs were identified based on insulation indices using TADtool (37). A/B compartmentalization was analyzed as previously described (38).

### Capture-C – Experimental procedure

Capture-C data were generated using the Next-Generation Capture-C protocol (34, 39). The DpnII restriction enzyme was used for digestion during 3C library preparation.

Because exclusion zones around all viewpoints analyzed in a multiplexed experiment are removed from analysis, we performed several Capture-C experiments to characterize the complete interaction landscapes of all *cis*-regulatory elements of interest in the globin loci. We used the following combinations of viewpoints in six independent experiments:

i. α-globin locus: R1 and R2 (enhancers); β-globin locus: HS1 and HS2 (enhancers);
ii. α-globin locus: R3, Rm and R4 (enhancers); β-globin locus: HS3 and HS4 (enhancers);
iii. α-globin locus: *Hbq-1* and *Hbq-2* (θ-globin promoters/CTCF-binding sites); β-globin locus: HS5 and HS6 (enhancers/CTCF-binding sites);
iv. α-globin locus: HS-38 and HS-39 (upstream CTCF-binding sites); β-globin locus: HS-57 (enhancer), HS-60 and HS-90 (upstream CTCF-binding sites);
v. α-globin locus: HS+44 and HS+48 (downstream CTCF-binding sites); β-globin locus: 3’HS1 and 3’HS2 (downstream CTCF-binding sites);
vi. α-globin locus: *Il9r, Snrnp25, Rhbdf1, Mpg* and *Nprl3* (promoters);

To cover the remaining regions of the α- and β-globin loci for the generation of a high-resolution all vs all contact matrix, we performed an additional multiplexed experiment with viewpoints tiled across a 300 kb window around both globin clusters.

Capture oligonucleotides were designed using CapSequm (34). Overviews of the viewpoint DpnII fragments in the α- and β-globin loci are shown in Supplementary Tables 1 and 2, respectively.

We used three biological replicates of primary murine ter119+ erythroid and ES cells in every experiment, which were pooled after ligation of indexed sequencing adapters.

The generated Capture-C libraries were sequenced using Illumina sequencing platforms (V2 chemistry; 150 bp paired-end reads).

### Capture-C – Data analysis

Capture-C data were analyzed as previously described (34). Reads were aligned to the mm9 reference genome using Bowtie1 with the following options: -p 1 -m 2 -v 3 –best –strata. The -m 2 option was used to allow for multi-mapping in the duplicated regions in the globin loci. The -v 3 option was used to allow up to three mismatches to improve alignment of the ES data in the β-globin region, which contains many SNPs in the 129/Ola strain compared to the mm9 reference. Because some SNPs were located in close proximity to or even in DpnII cut sites, alignment of reads with viewpoints in these DpnII fragments remained sub-optimal. The quality of some Capture-C profiles in the β-globin region in ES cells is therefore somewhat compromised, though still highly interpretable.

As PCR duplicates are removed during data analysis, Capture-C accurately quantifies chromatin interactions (40, 41). The Capture-C profiles in the figures represent the mean number of unique interactions per restriction fragment from three replicates, normalized for a total of 100,000 interactions on the chromosome analyzed, and scaled to 1,000.

### Tri-C – Experimental procedure

The R2 enhancer and HS-39 CTCF-binding site in the α-globin locus, and the HS2 enhancer and 3’HS1 CTCF-binding site in the β-globin locus are located on small NlaIII restriction fragment (Supplementary Table 3). We therefore used the NlaIII restriction enzyme for digestion of 3C libraries, which were prepared as previously described (34). To be able to capture multiple restriction fragments in individual reads, we optimized the subsequent sonication step to generate DNA fragments of ∼450 bp, using a Covaris S220 Focussed Ultra-Sonicator with the following settings: one cycle of 55 s; duty cycle: 10%; intensity: 4; cycles per burst: 200. DNA clean-up and size selection after sonication were performed with Ampure XP beads in a 0.7:1 bead-sample ratio. Subsequent ligation of sequencing adapters and indexing and multiplexing of libraries were performed as described previously (34). To enrich for reads containing the viewpoint fragments of interest, we performed a double oligonucleotide capture as previously described (34), using a 13 fmol pool of capture oligonucleotides.

We initially performed an experiment with three biological replicates of primary erythroid cells, which was sequenced using the Illumina MiSeq platform (V2 chemistry; 250 bp paired-end reads). *In silico* trimming of the 250 bp paired-end reads showed that there was little benefit of using 500 cycles of sequencing compared to 300 cycles for capturing multiple reporters in the reads. Therefore, to generate data at sufficient depth, we performed a second multiplexed experiment with seven additional technical replicates (derived from three biological replicates) of erythroid and ES cells, which was sequenced using the Illumina NextSeq platform (V2 chemistry; 150 bp paired-end reads).

### Tri-C – Data analysis

Tri-C data were analyzed using a combination of publicly available tools and customized scripts. Trim_galore (Babraham Institute, http://www.bioinformatics.babraham.ac.uk/projects/trim_galore/) was used to remove adapter sequences in the reads. Where possible, paired-end reads were reconstructed into single reads using FLASH with interleaved output settings. A custom script was used to perform an *in silico* restriction enzyme digestion, after which the reads were aligned to the reference genome using Bowtie1 (-p 1 -m 2 -v 3 –best –strata). The aligned reads were analyzed with custom scripts to identify captured reads containing the targeted viewpoints of interest. Restriction fragments in captured reads were defined as interacting ‘reporter’ fragments if they were located outside a ∼1 kb exclusion zone around the viewpoint restriction fragment. PCR duplicates were removed by excluding reads that had the same start and end coordinates of each individual restriction fragment. Reads with two or more reporters were used to calculate interaction frequencies between reporter fragments for each viewpoint. An overview of the numbers of detected reporters is shown in Supplementary Table 4. The interaction frequencies were normalized for a total of 100,000 interactions on the chromosome analyzed and binned in 500 bp bins, to allow the data to be represented in symmetrical contact matrices.

### C-Trap – Experimental procedure

3C libraries were prepared as previously described (34), using the DpnII restriction enzyme for digestion. Primers targeting the restriction fragments of interest were designed >100 bp away from the restriction site (Supplementary Table 5), to allow for the selection of reads that were selectively amplified and of sufficient quality. PCR amplification was performed using the Takara Prime Star GXL 2-step amplification program for 10-30 kb amplicons. The Agencourt AMPure XP system was used to purify the PCR product and select fragments >600 bp. The purified amplicons were adapter-ligated and sequenced using the Oxford Nanopore MinION platform (MAP-005 chemistry).

### C-Trap – Data analysis

The MinION reads were basecalled using Metrichor and converted to fasta format using Poretools (43). To select reads that were specifically amplified and of sufficient quality, a BLAST search against a database containing the sequences between the primers and the restriction sites of the targeted fragments was performed. Reads that matched >60% of both primer sequences were selected. To map the trapped restriction fragments in the reads, a BLAST search against a database containing all the DpnII fragments in the genome was used. Custom scripts were used to iterate through the BLAST results and select the non-overlapping BLAST matches in the reads with a match >60% of fragment length or >100 bp (to avoid a skew towards smaller fragments) in order of significance of the BLAST scores. A summary of the read statistics is shown in Supplementary Table 6.

---

**Supplementary Figure 1:**
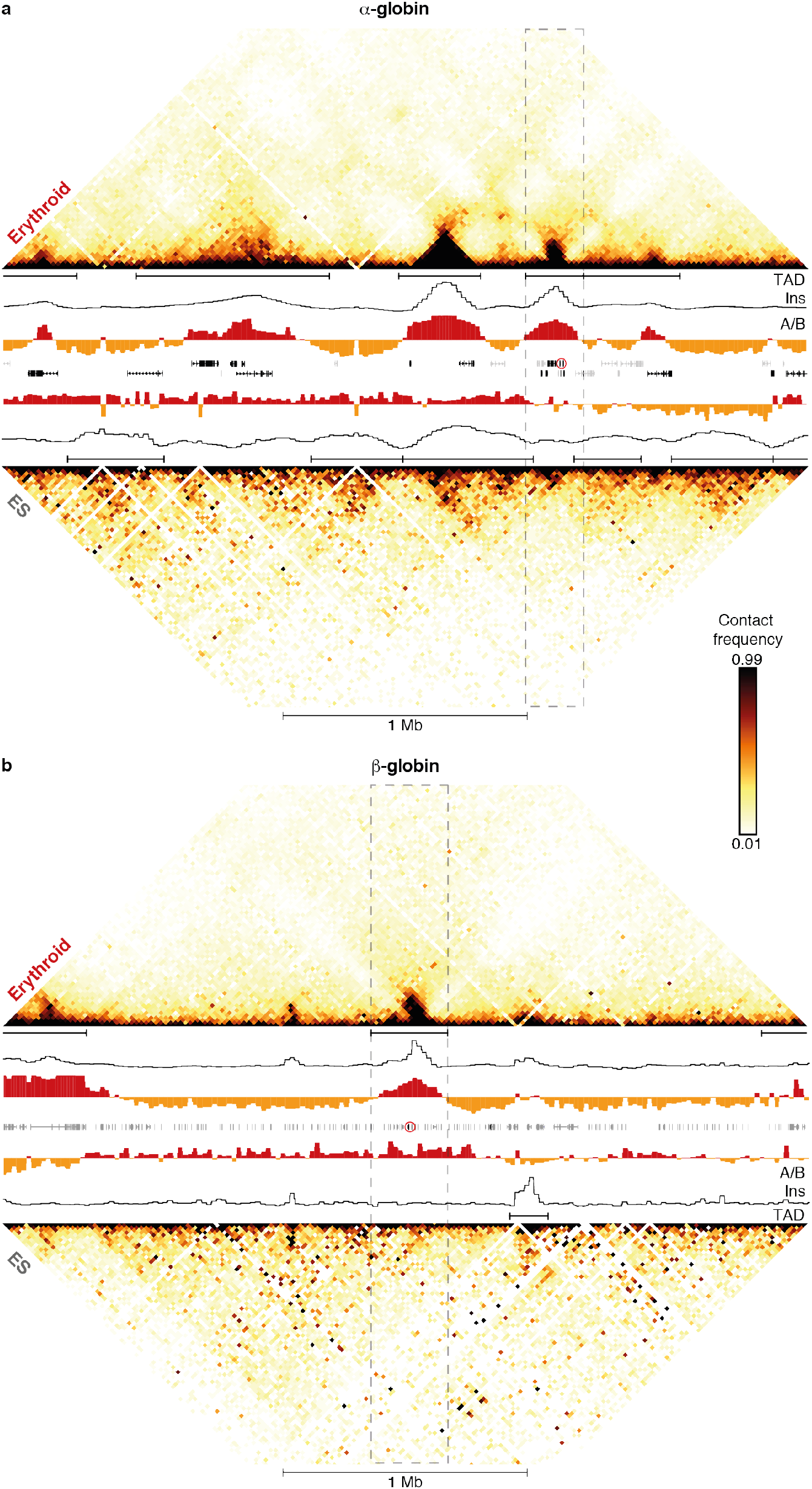
The organization of the extended globin loci into self-interacting domains is tissue-specific. Hi-C contact matrices (20 kb resolution) of the **(a)** α-globin and the **(b)** β-globin loci in erythroid (top) and ES (bottom) cells. Topologically Associating Domains (TAD), insulation indices (Ins), A/B-compartmentalization (A/B; red/yellow), and gene annotation (active genes in black, inactive genes in grey, globin genes highlighted in red) are shown in the middle and the TADs containing the globin loci are highlighted in dashed grey boxes. Coordinates (mm9): chr11:29,900,000-33,200,000 (α-globin) and chr7:109,300,000-112,600,000 (β-globin).

**Supplementary Figure 2:**
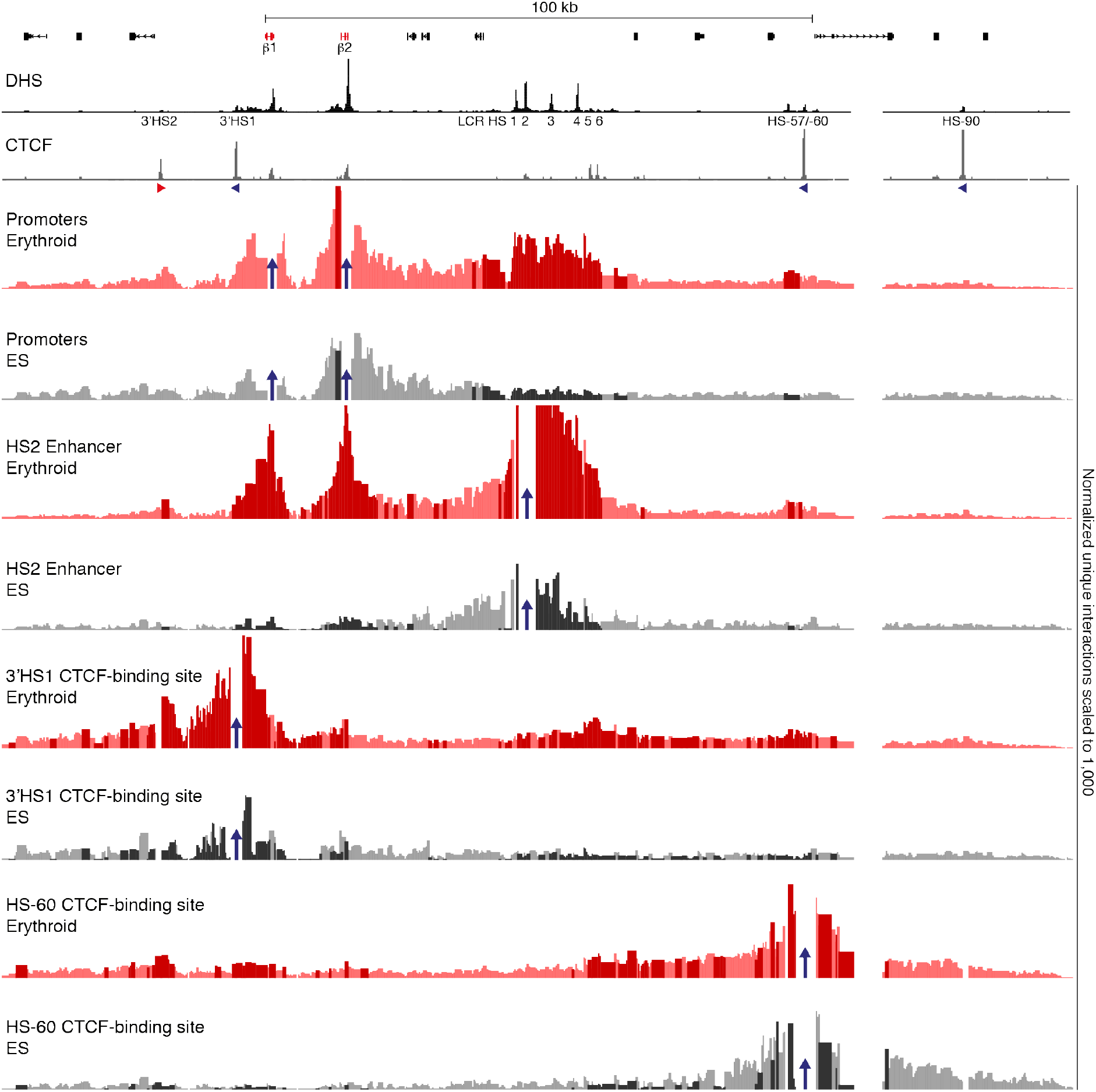
Characterization of the interaction landscape of the regulatory elements of the β-globin locus. High-resolution Capture-C interaction profiles of the β-globin locus from the viewpoints (indicated by blue arrows) of the β-globin promoters, the HS2 enhancer, and CTCF-binding sites 3’HS1 and HS-60 in erythroid (red) and ES (grey) cells. Profiles represent the mean number of normalized unique interactions per restriction fragment from three replicates. Significantly different interactions between erythroid and ES cells are highlighted in bold colors. Gene annotation (β-globin genes highlighted in red), erythroid DNaseI Hypersensitive Sites (DHS) and CTCF occupancy are shown at the top, with arrows indicating the orientation of the CTCF-binding motifs. Coordinates (mm9): chr7:110,912,000-111,112,000.

**Supplementary Figure 3:**
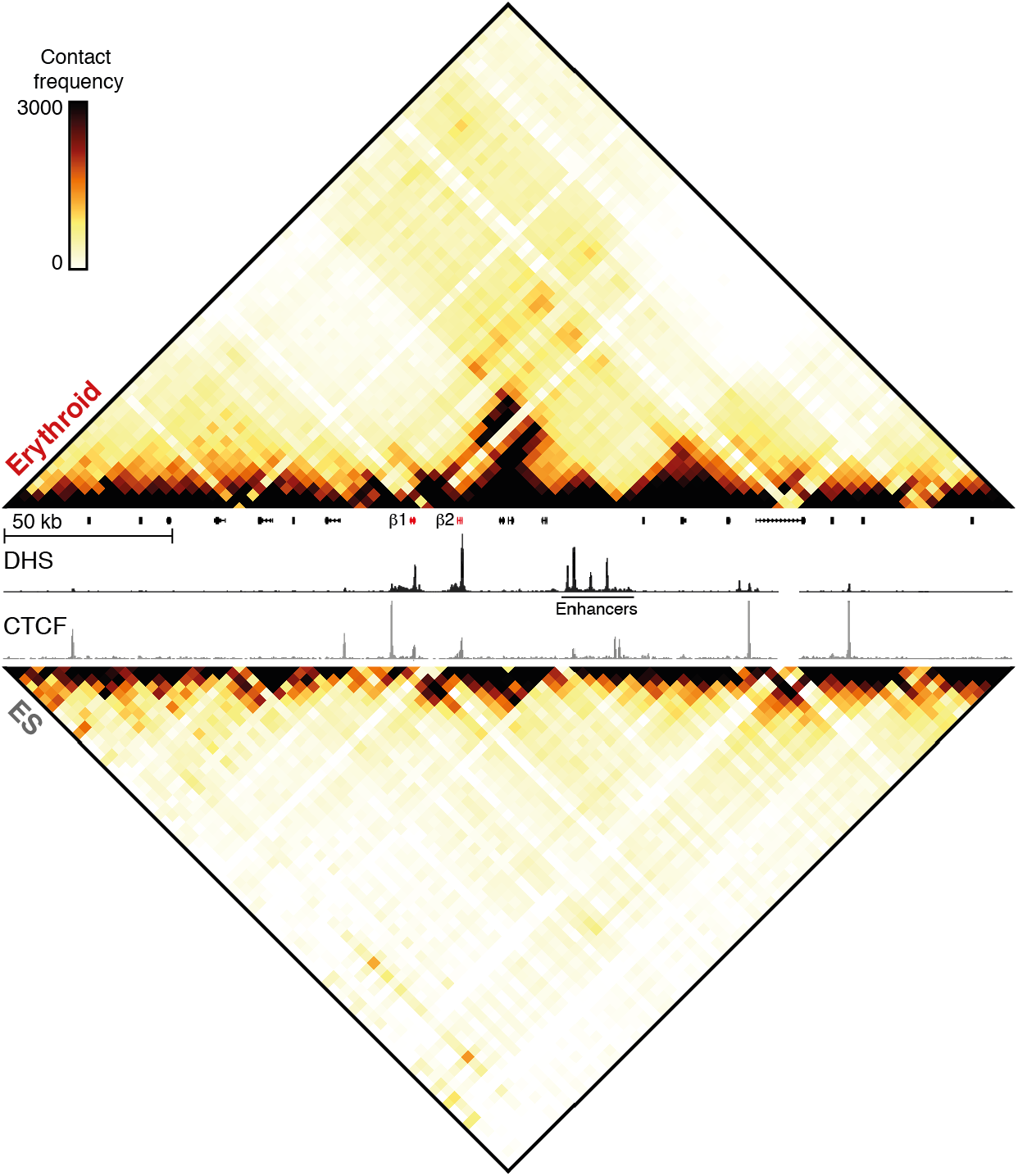
Structural conformation of the active and inactive β-globin locus. Contact matrices (4 kb resolution) of the β-globin locus derived from Capture-C experiments with viewpoints closely spaced across the domain in erythroid (top) and ES (bottom) cells. Contact frequencies represent the mean number of normalized unique interactions from three replicates. Gene annotation (β-globin genes highlighted in red), erythroid DNaseI Hypersensitive Sites (DHS) and CTCF occupancy are shown in the middle. Coordinates (mm9): chr7:110,840,000-111,140,000.

**Supplementary Figure 4:**
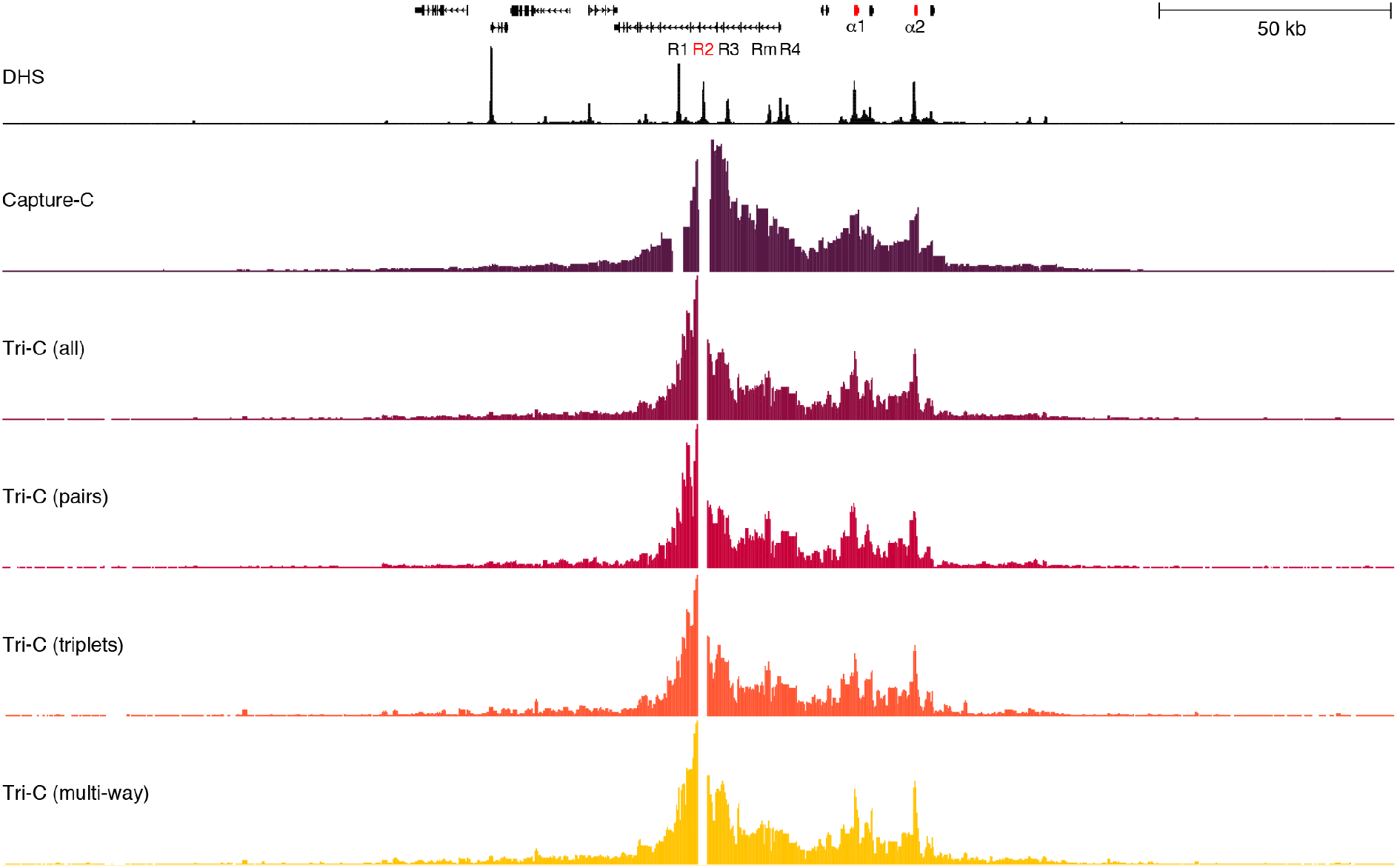
Validation of pair-wise interactions detected by Tri-C. Comparison of the Capture-C interaction profile (purple) from the viewpoint of the R2 enhancer in the α-globin locus to pair-wise interactions derived from Tri-C data. The first Tri-C profile (maroon) shows pair-wise interactions derived from all unique Tri-C reads; the tracks below compare pair-wise interactions detected in unique Tri-C reads in which two (red), three (orange) or multiple (yellow) interacting fragments were detected. The interaction profiles are very similar, demonstrating that detection of multiple ligation events simultaneously does not skew the data. The subtle differences between the Capture-C and Tri-C profiles are likely related to the use of different restriction enzymes during 3C library preparation (DpnII in Capture-C; NlaIII in Tri-C). Gene annotation (α-globin genes highlighted in red) and erythroid DNaseI Hypersensitive Sites (DHS) are shown at the top. Coordinates (mm9): chr11:32,000,000-32,300,000.

**Supplementary Figure 5:**
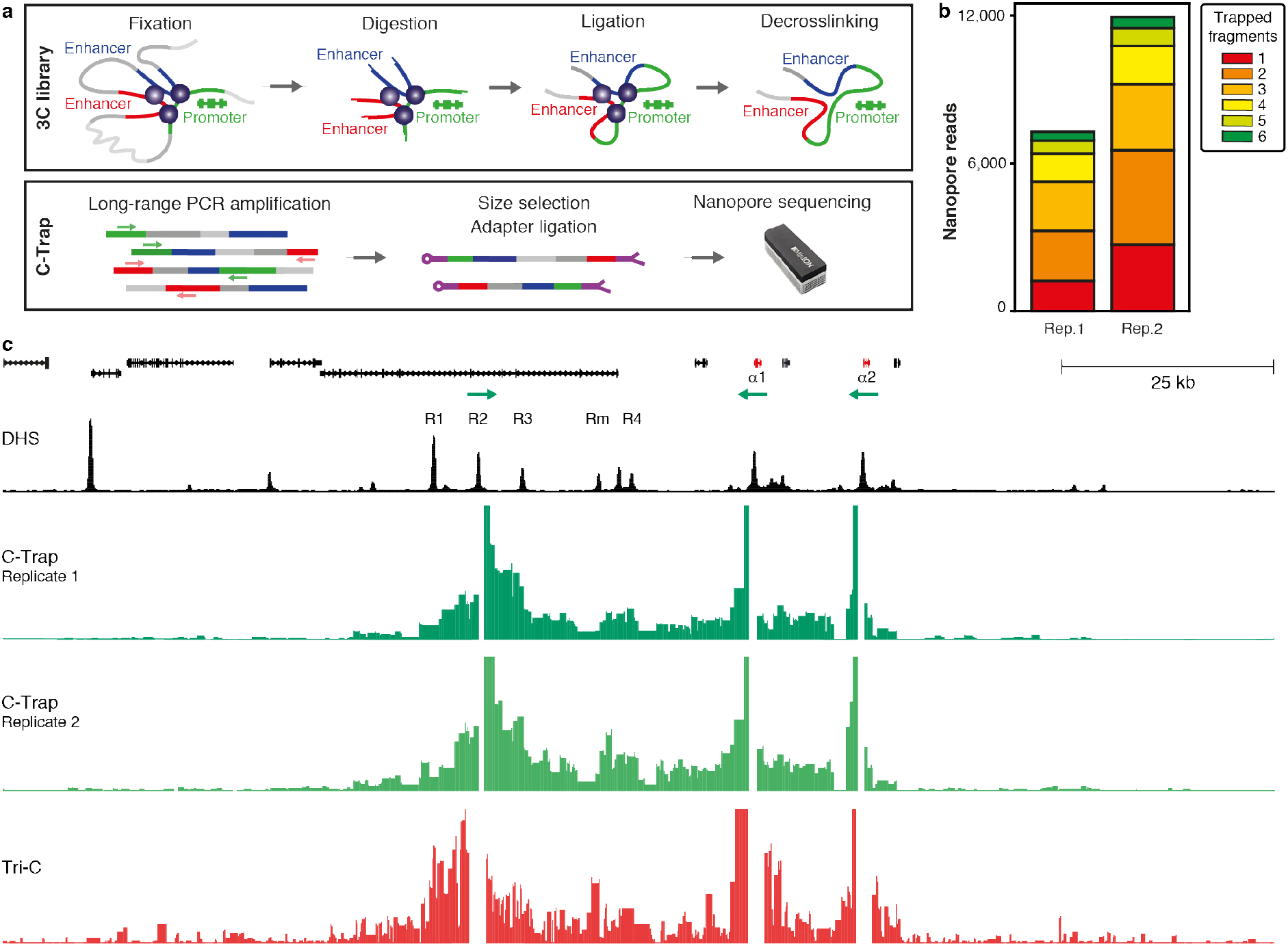
Validation of multi-way interactions detected by Tri-C. **(a)** C-Trap is a novel 3C approach that can analyze interactions occurring simultaneously with an interaction between two fragments of interest. It uses long-range PCR amplification with primers targeting these two fragments to enrich for ligation products in the 3C library that contain the interaction of interest and ‘trap’ the intervening fragments that were interacting simultaneously. After size selection to deplete PCR products <700bp, the Nanopore MinION long-read sequencing platform is used to detect all trapped fragments. **(b)** Overview of the number of trapped fragments per read detected in a C-Trap experiment in which the interaction between the R2 enhancer and the α-globin promoters was targeted in erythroid cells (two biological replicates). Read numbers represent reads in which both primer sequences could be detected at the ends. These numbers are much lower compared to a Tri-C experiment (Figure 3b), due to the low output and read quality of the Nanopore MinION sequencing platform. The sensitivity of C-Trap is therefore ∼100-fold lower compared to Tri-C. Though C-Trap has the advantage of being capable of identifying many simultaneously interacting fragments in ligation products, the majority of the reads contain only a few trapped fragments. This could reflect the strength of the interaction between the R2 enhancer and the α-globin promoters, which are therefore often in close proximity in 3C ligation products, with few intervening fragments. It might also be related to a PCR amplification skew towards preferential enrichment of shorter fragments and lower quality of longer Nanopore reads. **(c)** Comparison of multi-way interaction profiles generated from C-Trap reads containing interacting fragments with the R2 enhancer and the α-globin promoters, and from Tri-C reads containing both the R2 viewpoint and the α-globin promoters, in primary erythroid cells. The profiles look very similar: interactions are confined to the strongly compartmentalized domain and enriched over the enhancers. This confirms the validity of the multi-way interactions detected by Tri-C. Gene annotation (α-globin genes highlighted in red) and erythroid DNaseI Hypersensitive Sites (DHS) are shown at the top. The location of the C-Trap primers is indicated by green arrows. Coordinates (mm9): chr11:32,095,000-32,245,000.

**Supplementary Figure 6:**
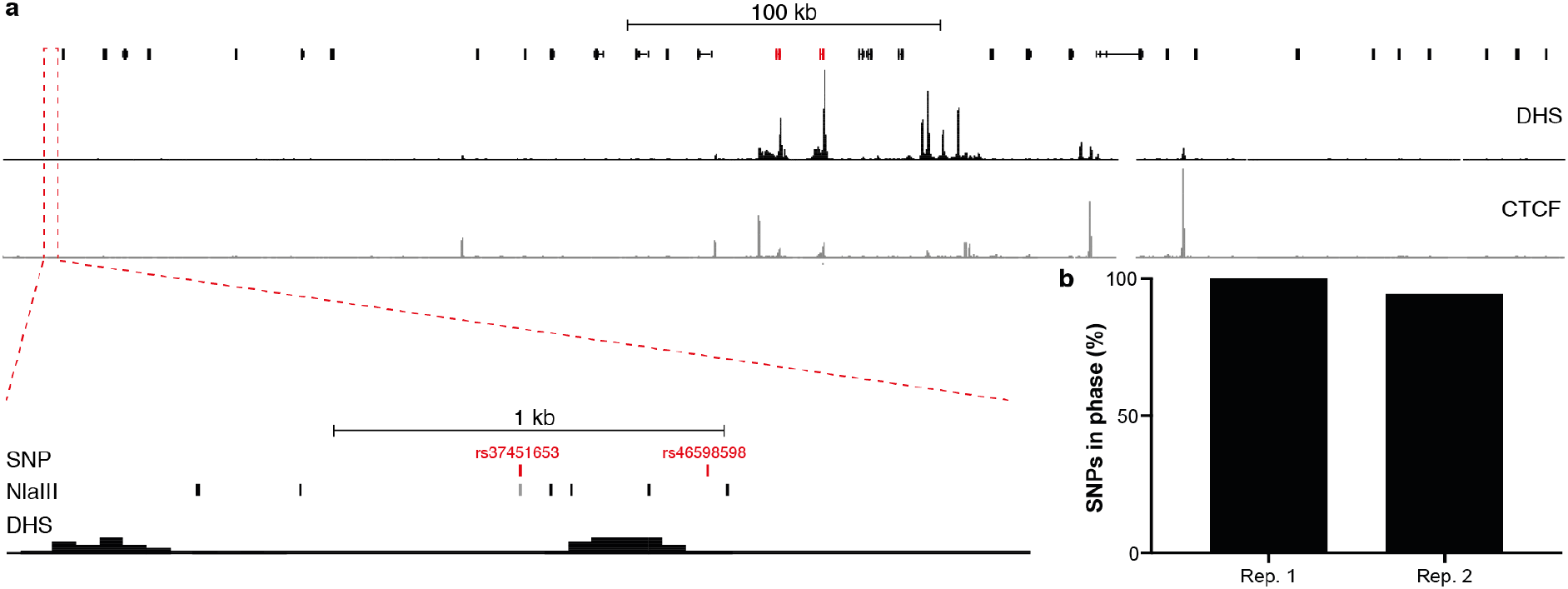
Interactions detected by Tri-C are allele-specific. By analyzing the distribution of heterozygous SNPs in F1 hybrids, we have previously shown that interactions detected by Capture-C predominantly (>95%) originate from the same allele (34). To confirm that multi-way interactions identified by Tri-C are also allele-specific, we analyzed the distribution of SNPs in the β-globin locus in ES cells. **(a)** E14 ES cells are heterozygous for SNPs rs37451653 (C/T) and rs46598598 (G/C), which are located downstream of the β-globin genes. The minor variant of rs37451653 creates a new NlaIII cut site (CACG/CATG). Gene annotation (β-globin genes highlighted in red), erythroid DNaseI Hypersensitive Sites (DHS) and CTCF occupancy are shown at the top. Coordinates (mm9): chr7:110,713,500-111,213,500. (b) The small NlaIII fragment created by the minor rs37451653 variant (T) is predominantly captured with the minor rs46598598 variant (C) in unique reads containing both SNPs in two independent ES replicates.

**Supplementary Figure 7:**
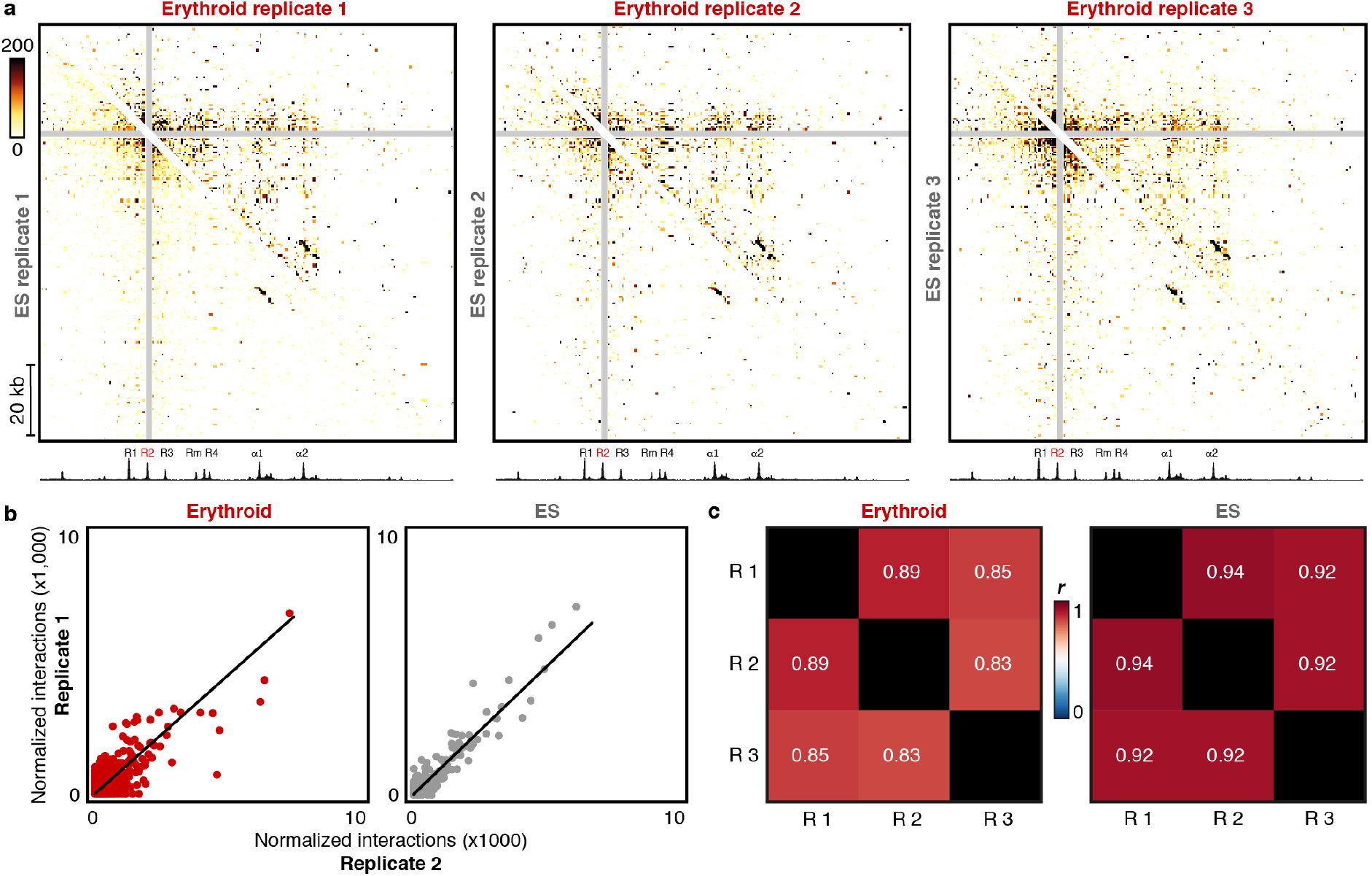
Tri-C data are highly reproducible. **(a)** Comparison of Tri-C contact matrices (500 bp resolution) from the viewpoint of the R2 enhancer in the α-globin domain in three biological replicates. Erythroid samples are shown in the right top half of the matrix and ES samples in the left bottom half. Proximity contacts around R2 are excluded (grey diagonal). Erythroid DNaseI Hypersensitive Sites are shown at the bottom of the matrices. Coordinates (mm9): chr11:32,120,000-32,240,000. **(b)** Correlations of multi-way interaction frequencies detected from the R2 viewpoint between individual erythroid (left) and ES (right) replicates. **(c)** Correlation matrices showing Pearson correlation coefficients (*r*) of multi-way interaction frequencies detected from the R2 viewpoint between replicates (R) of erythroid (left) and ES (right) cells.

**Supplementary Figure 8:**
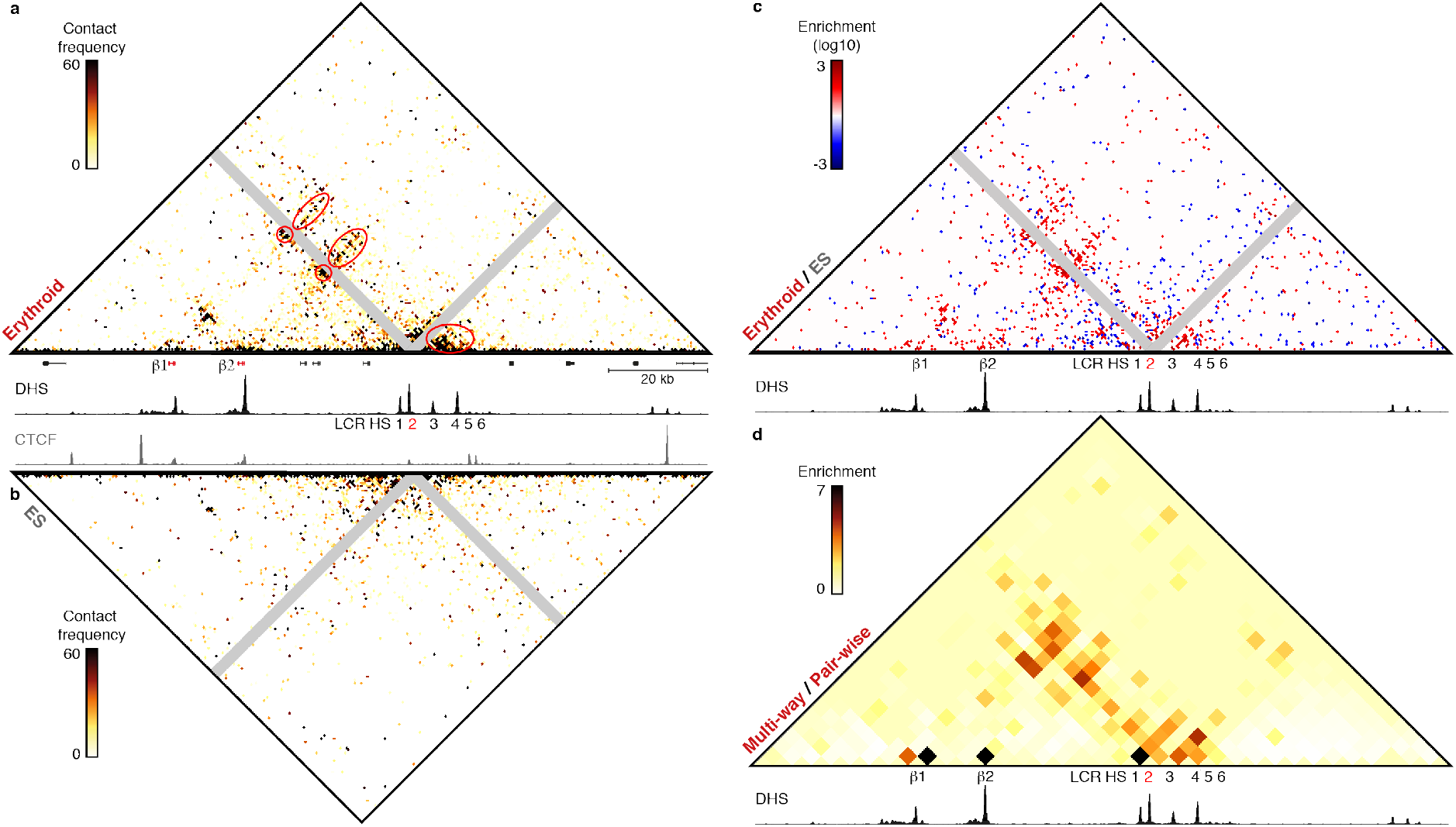
Analysis of multi-way interactions between enhancers and promoters in the β-globin locus. Tri-C data (500 bp resolution) showing multi-way chromatin interactions with the HS2 enhancer in the β-globin locus. Gene annotation (β-globin genes highlighted in red), erythroid DNaseI Hypersensitive Sites (DHS) and/or CTCF-binding sites are shown at the bottom of the matrices, **(a)** Contact matrix showing normalized multi-way interactions in three replicates of erythroid cells. Proximity contacts around HS2 are excluded (grey diagonal) and specific enrichments are highlighted (red circles/ovals). **(b)** Contact matrix showing normalized multi-way interactions in three replicates of ES cells. Proximity contacts around HS2 are excluded (grey diagonal). **(c)** Contact matrix highlighting interactions that are >20-fold enriched in erythroid (red) or ES (blue) cells. Proximity contacts around HS2 are excluded (grey diagonal). **(d)** Contact matrix (4 kb resolution) showing multi-way interactions after correcting for the pair-wise contact frequencies derived from the multiplexed Capture-C data. Coordinates (mm9): chr7:110,930,000-111,070,000.

**Supplementary Figure 9:**
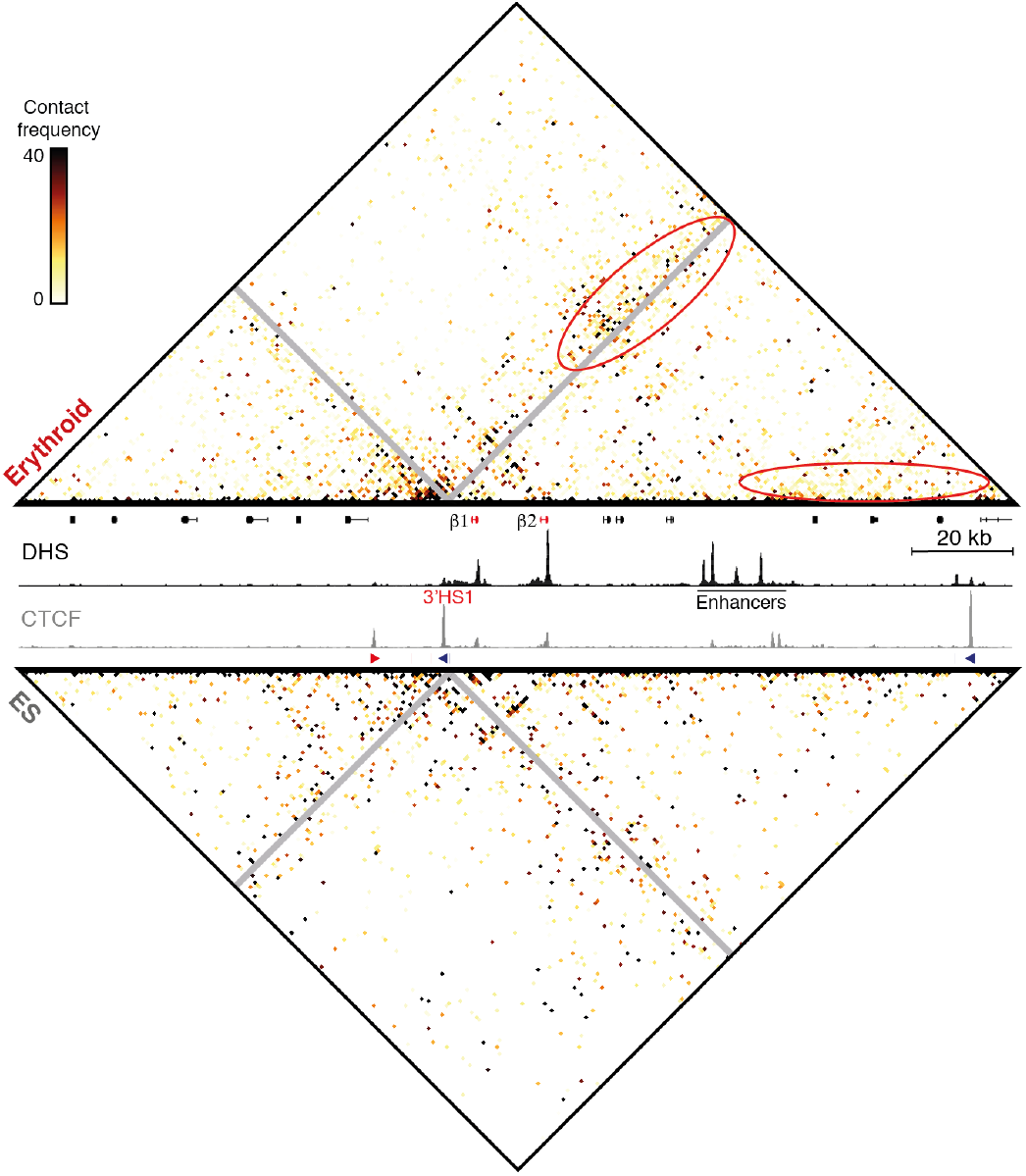
Analysis of multi-way interactions between CTCF-binding sites in the β-globin locus. Tri-C contact matrices (500 bp resolution) showing normalized multi-way chromatin interactions with CTCF-binding site 3’HS1 in the β-globin locus in three replicates of erythroid (top) and ES (bottom) cells. Gene annotation (β-globin genes highlighted in red), erythroid DNaseI Hypersensitive Sites (DHS) and CTCF-binding sites are shown in the middle, with arrows indicating the orientation of the CTCF-binding motifs. Proximity contacts around 3’HS1 are excluded (grey diagonal) and specific enrichments in erythroid cells are highlighted (red ovals). Coordinates (mm9): chr7:110,870,000-111,070,000.

**Supplementary Table 1:** Capture-C oligonucleotides targeting the α-globin locus. Overview of the viewpoints analyzed in Capture-C experiments in the α-globin locus. Experiments were performed with two 120 bp capture oligonucleotides targeting both ends of the viewpoint DpnII fragments. When one of the ends contained a repetitive sequence, only one oligonucleotide targeting the unique end was used. The table shows that 120 bp end of the restriction fragment instead of the entire fragment in that case.

| Viewpoint | Annotation | DpnII fragment |
| --- | --- | --- |
| Hba-1 | Promoter | chr11:32182969-32183821 |
| Hba-2 | Promoter | chr11:32195804-32196638 |
| R1 | Enhancer | chr11:32145277-32146568 |
| R2 | Enhancer | chr11:32151060-32151883 |
| R3 | Enhancer | chr11:32155865-32156383 |
| Rm | Enhancer | chr11:32164966-32165085 |
| R4 | Enhancer | chr11:32168680-32169413 |
| Hbq-1 | Promoter/CTCF-binding site | chr11:32199491-32200139 |
| Hbq-2 | Promoter/CTCF-binding site | chr11:32186467-32187034 |
| HS-38 | CTCF-binding site | chr11:32138079-32139000 |
| HS-39 | CTCF-binding site | chr11:32137176-32137426 |
| HS+44 | CTCF-binding site | chr11:32220918-32221720 |
| HS+48 | CTCF-binding site | chr11:32224323-32227298 |
| II9r | Promoter | chr11:32099819-32100299 |
| Snrnp25 | Promoter | chr11:32104921-32105535 |
| Rhbdf1 | Promoter | chr11:32122089-32122693 |
| Mpg | Promoter | chr11:32126341-32126674 |
| Nprl3 | Promoter | chr11:32167508-32168372 |
| 32003k | Model_viewpoint | chr11:32002284-32002785 |
| 32012k | Model_viewpoint | chr11:32011827-32012022 |
| 32019k | Model_viewpoint | chr11:32019083-32019233 |
| 32027k | Model_viewpoint | chr11:32027129-32027804 |
| 32037k | Model_viewpoint | chr11:32036007-32037157 |
| 32042k | Model_viewpoint | chr11:32042139-32042440 |
| 32051k | Model_viewpoint | chr11:32050692-32051800 |
| 32059k | Model_viewpoint | chr11:32059255-32059583 |
| 32068k | Model_viewpoint | chr11:32067632-32068398 |
| 32077k | Model_viewpoint | chr11:32076914-32077034 |
| 32084k | Model_viewpoint | chr11:32083717-32083891 |
| 32091k | Model_viewpoint | chr11:32090436-32092064 |
| 32111k | Model_viewpoint | chr11:32110960-32111206 |
| 32119k | Model_viewpoint | chr11:32118902-32119562 |
| 32175k | Model_viewpoint | chr11:32175103-32175364 |
| 32207k | Model_viewpoint | chr11:32207138-32207281 |
| 32216k | Model_viewpoint | chr11:32215331-32215799 |
| 32236k | Model_viewpoint | chr11:32235766-32237204 |
| 32243k | Model_viewpoint | chr11:32242417-32243041 |
| 32253k | Model_viewpoint | chr11:32252649-32252858 |
| 32259k | Model_viewpoint | chr11:32258095-32259644 |
| 32267k | Model_viewpoint | chr11:32267139-32267638 |
| 32275k | Model_viewpoint | chr11:32274955-32275418 |
| 32284k | Model_viewpoint | chr11:32283727-32283847 |
| 32291k | Model_viewpoint | chr11:32291086-32291835 |
| 32298k | Model_viewpoint | chr11:32297594-32298445 |

**Supplementary Table 2:** Capture-C oligonucleotides targeting the β-globin locus. Overview of the viewpoints analyzed in Capture-C experiments in the β-globin locus. Experiments were performed with two 120 bp capture oligonucleotides targeting both ends of the viewpoint DpnII fragments. When one of the ends contained a repetitive sequence, only one oligonucleotide targeting the unique end was used. The table shows that 120 bp end of the restriction fragment instead of the entire fragment in that case.

| Viewpoint | Annotation | DpnII fragment |
| --- | --- | --- |
| Hbb-1 | Promoter | chr7:110961967-110962817 |
| Hbb-2 | Promoter | chr7:110975976-110976456 |
| HS1 | Enhancer (LCR) | chr7:111007506-111007930 |
| HS2 | Enhancer (LCR) | chr7:111009550-111009749 |
| HS3 | Enhancer (LCR) | chr7:111014164-111014602 |
| HS4 | Enhancer (LCR) | chr7:111018972-111019398 |
| HS5 | Enhancer (LCR) | chr7:111021350-111022005 |
| HS6 | Enhancer (LCR) | chr7:111023278-111023655 |
| 3'HS1 | CTCF-binding site | chr7:110955508-110955666 |
| 3'HS2 | CTCF-binding site | chr7:110940624-110941597 |
| HS-57 | Enhancer | chr7:111058110-111058705 |
| HS-60 | CTCF-binding site | chr7:111061643-111062050 |
| HS-90 | CTCF-binding site | chr7:111091378-111092626 |
| 110846k | Model_viewpoint | chr7:110845602-110845722 |
| 110852k | Model_viewpoint | chr7:110851723-110852505 |
| 110863k | Model_viewpoint | chr7:110862597-110862717 |
| 110870k | Model_viewpoint | chr7:110869104-110870170 |
| 110879k | Model_viewpoint | chr7:110879128-110879377 |
| 110887k | Model_viewpoint | chr7:110885985-110887299 |
| 110894k | Model_viewpoint | chr7:110894213-110894515 |
| 110902k | Model_viewpoint | chr7:110901673-110903198 |
| 110909k | Model_viewpoint | chr7:110908816-110908936 |
| 110919k | Model_viewpoint | chr7:110918844-110919286 |
| 110928k | Model_viewpoint | chr7:110926852-110928205 |
| 110934k | Model_viewpoint | chr7:110934018-110934138 |
| 110971k | Model_viewpoint | chr7:110971166-110971561 |
| 110983k | Model_viewpoint | chr7:110983084-110983538 |
| 110991k | Model_viewpoint | chr7:110990420-110991222 |
| 111000k | Model_viewpoint | chr7:110999928-111000287 |
| 111027k | Model_viewpoint | chr7:111026798-111027050 |
| 111036k | Model_viewpoint | chr7:111036248-111036368 |
| 111043k | Model_viewpoint | chr7:111043069-111043411 |
| 111051k | Model_viewpoint | chr7:111051223-111051670 |
| 111068k | Model_viewpoint | chr7:111068238-111068358 |
| 111078k | Model_viewpoint | chr7:111078043-111078649 |
| 111087k | Model_viewpoint | chr7:111086801-111087189 |
| 111100k | Model_viewpoint | chr7:111100104-111100247 |
| 111106k | Model_viewpoint | chr7:111106311-111106563 |
| 111122k | Model_viewpoint | chr7:111121964-111122084 |
| 111129k | Model_viewpoint | chr7:111129322-111129442 |
| 111139k | Model_viewpoint | chr7:111138889-111139009 |

**Supplementary Table 3:** Tri-C capture oligonucleotides. Overview of the viewpoints analyzed with Tri-C. Experiments were performed with 120 bp capture oligonucleotides targeting the middle of the viewpoint NlaIII fragments.

| Viewpoint | Annotation | Capture oligonucleotide | NlaIII fragment size (bp) |
| --- | --- | --- | --- |
| R2 | Enhancer (α-globin) | chr11:32150965-32151084 | 176 |
| HS-39 | CTCF-binding site (α-globin) | chr11:32137199-32137318 | 136 |
| HS2 | Enhancer (β-globin) | chr7:111009445-111009564 | 216 |
| 3'HS1 | CTCF-binding site (β-globin) | chr7:110955472-110955591 | 143 |

**Supplementary Table 4:** Tri-C read statistics. Overview of the number of reads detected in erythroid and ES samples in the multiplexed Tri-C experiments. Numbers represent unique reads after PCR duplicate removal. The first row shows the total number of reads containing both the viewpoint and at least one reporter fragment. These are subdivided in reads containing one, two or more reporters in the rows below. The bottom row shows the total number of reporters detected.

|  | Erythroid |  |  |  | ES |  |  |  |
| --- | --- | --- | --- | --- | --- | --- | --- | --- |
| | $\alpha$: R2 | $\alpha$: HS-39 | $\beta$: HS2 | $\beta$: 3'HS1 | $\alpha$: R2 | $\alpha$: HS-39 | $\beta$: HS2 | $\beta$: 3'HS1 |
| Reads (total) | 1,543,888 | 1,382,231 | 1,637,730 | 3,313,613 | 1,500,991 | 1,026,015 | 949,036 | 1,398,212 |
| - 1 reporter | 806,832 | 673,174 | 859,769 | 1,555,070 | 847,963 | 492,204 | 531,019 | 652,602 |
| - 2 reporters | 514,710 | 479,530 | 550,410 | 1,149,968 | 473,430 | 369,812 | 309,108 | 505,782 |
| - >2 reporters | 222,346 | 229,527 | 227,551 | 608,575 | 179,598 | 163,999 | 108,909 | 239,828 |
| Reporters (total) | 2,555,425 | 2,379,973 | 2,694,822 | 5,847,628 | 2,367,219 | 1,757,001 | 1,495,350 | 2,437,140 |

**Supplementary Table 5:**
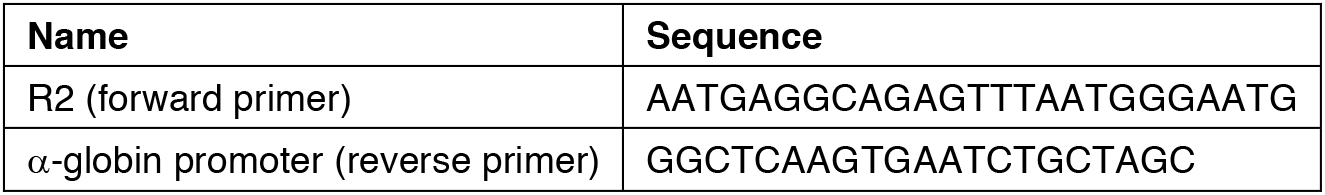
C-Trap primers. Overview of the primers used for PCR enrichment in the C-Trap experiments targeting the interaction between the R2 enhancer and the α-globin promoter.

**Supplementary Table 6:**
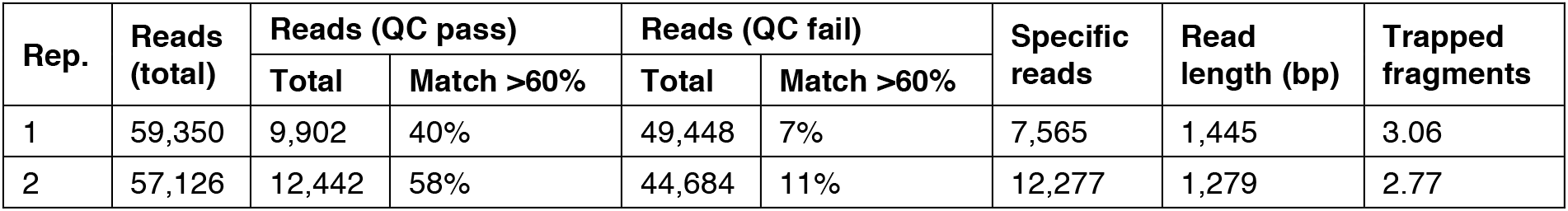
C-Trap read statistics. Overview of the number of reads detected in C-Trap experiments in two replicates (rep.) of primary erythroid cells. The first column shows the number of basecalled reads in each replicate. Approximately half of the reads that passed the Metrichor quality control (QC) matched both primer sequences; these were defined as specific reads and used for analysis. Of the reads that failed QC, only a small percentage could be used for further analysis. The last two columns show the average read length and number of trapped fragments in the specific reads detected.

